# Use of Protein interactions from Imaging Complexes after Translocation (PICT) to characterise *in situ* the spatial configuration of proteins interacting with the exocyst

**DOI:** 10.1101/2024.03.28.587178

**Authors:** Altair C. Hernandez, Laura I. Betancur, Andrea Picco, Oriol Gallego

**Author notes:** Altair C. Hernandez and Laura I. Betancur have equally contributed to this work.

## Abstract

Although the structure of the exocyst has been successfully resolved by cryo-electron microscopy, multiple studies showed that exocyst function requires the transient interaction with additional proteins. Unfortunately, the exocyst-interacting network could not be collectively reconstituted, challenging the understanding of how the exocyst complex is coordinated within the network of proteins involved in exocytosis. In a previous work, we described an approach that combines Protein interactions from Imaging Complexes after Translocation (PICT) and centroid localization analysis of diffraction-limited fluorescence signals to estimate the distance between a labelled protein and a spatial reference. This approach allows resolving the spatial organisation of protein interactions directly in living cells, both for intra-complex (i.e. between exocyst subunits) and inter-complex (i.e. between exocyst and transient binding proteins) interactions. In this chapter, we present the protocol to reproduce the sample preparation and image acquisition for PICT experiments. We also describe the computational image analysis pipeline to estimate the distance in PICT experiments. As illustration of the approach, we measure the distance from the spatial reference where the exocyst is anchored to 1) an intra-complex interaction (i.e. Sec5 exocyst subunit) and 2) an inter-complex interaction (Sec2, a guanyl-nucleotide exchange factor mediating vesicle tethering).

## 1. Introduction

Fluorescence microscopy allows investigating the spatial organisation of biological macromolecules *in vivo*, providing a unique opportunity to tackle the structural analysis of protein complexes into its native environment. While light diffraction limits the resolution of fluorescence microscopy to 200–300 nm, localisation microscopy can estimate the centroid position of fluorescently labelled proteins with 20–30 nm precision (**1**). By localising diffraction-limited fluorescent spots, light microscopy enables the measurement of the separation between fluorescently labelled proteins. Reproducible measurements of the separation provide estimates with a precision of 1 nm (**2–6**). Unfortunately, resolving the 3D architecture of macromolecular complexes *in vivo* remains a challenge due to the intrinsic dynamism of protein networks, which hinders obtaining reproducible measurements necessary to achieve high precision in living cells.

PICT (Protein interactions from Imaging Complexes after Translocation) is a live-cell imaging method initially developed for the quantitative characterization of protein-protein interactions (**7**). PICT is based on the anchor-away method (**8**), which uses the rapamycin-induced heterodimerization of FK506-binding protein (FKBP) and FK506-rapamycin binding (FRB) domain (**9**). With PICT, rapamycin treatment induces the recruitment of a FRB-tagged bait protein (bait-FRB) to a static anchoring platform defined by the endocytic protein Sla2 fused to RFP and FKBP (Sla2-RFP-FKBP) (**10**). Proteins that directly or indirectly bind the bait-FRB are also co-recruited to the anchoring platform. This co-recruitment can be visualised as a change in the localisation of the interacting protein labelled with mNeonGreen (prey-mNG) which forms diffraction-limited spots co-localising with the Sla2-RFP-FKBP diffraction-limited spots. Thus, PICT enables the reproducible ectopic localization of the bait-FRB to the anchoring platform associated with the yeast plasma membrane. Static anchoring platforms allow for large exposure imaging, maximising the detection of photons emitted by the RFP and mNG fluorescent proteins and enabling the fluorophore-to-fluorophore distance measurement localisation of the centroid of fluorescence with 2–5 nm precision. The labelling of different proteins with mNG allows measuring the distance from each prey-mNG to the Sla2-RFP-FKBP that acts as a spatial reference.

The exocyst complex is a hetero-octamer composed of two subcomplexes (module-I: Sec3-Sec5-Sec6-Sec8, and module-II: Sec10-Sec15-Exo70-Exo84) (**11**). Exocyst function involves transient interactions with secretory vesicle-associated proteins (**12, 13**), plasma membrane phospholipids (PI(4,5)P_2_) (**14–16**), Sec1/Munc-18 proteins (**17, 18**), soluble *N-*ethylmaleimide-sensitive factor attachment protein receptor (SNARE) proteins (**16, 19–23**), small GTPases (**24–27**) and others. Some of these interactions have been suggested to induce exocyst conformational changes required for its function (**28, 29**). The structural analysis of the exocyst using x-ray crystallography and cryo-electron microscopy combined with chemical cross-linking and homology modelling, have elucidated almost 80% of the exocyst structure at near-atomic resolution (**11, 30**). However, the need for protein purification and the absence of physiological measurements in the *in vitro* reconstitution challenges the functional interpretation of the high-resolution model. To fully understand the exocyst mechanism of action, it is necessary to analyse its structure in its physiological context: the cell. In the past, we integrated PICT measurements together with available structural features to model the architecture of the exocyst complex in its vesicle-bound state in living cells (**10**). Completing the functional architecture of the exocyst together with its regulatory-interacting proteins will be pivotal for unravelling the molecular mechanisms that govern exocytosis.

In this chapter, we outline the protocol for conducting PICT experiments and estimating the distance between labelled prey proteins and Sla2-RFP-FKBP. As a proof-of-concept, we present the analysis of distances between Sla2-RFP-FKBP and the labelled prey proteins Sec5-mNG and Sec2-mNG when using Exo70-FRB as bait. The measurements suggest that multiple Sec2-mNG are evenly distributed throughout the vesicle surface when the vesicle is bound by the exocyst.

## 2. Materials

### 2.1 Equipment

● Epifluorescence microscope: We use an automated ECLIPSE Ti2-E inverted microscope (Nikon) equipped with a motorised stage, a 100x magnification and 1.49 numerical aperture (NA) SR HP APO TIRF objective (Nikon), and a Zyla 4.2 sCMOS camera (Andor) with 2048 x 2048 pixels and 6.5 µm pixel size. Fluorophores are excited with a light-emitting diode (LED) illumination system (SpectraX Lumencor). The excitation light is filtered with a 475/35-nm filter (Chroma) for mNG and a 575/35-nm filter (Chroma) for RFP. Channels are acquired sequentially using a FF505/606-Di01-25x36 dual band-pass dichroic mirror (Semrock) and the emission light is filtered with a FF01-524/628-25 dual band-pass filter (Semrock) (*see* **Note 1**). The entire system is controlled by µManager 2.0 software (**31**) and the microscope features a Perfect Focus System (PFS) (Nikon) to minimise axial drift.
● Bench centrifuge and microcentrifuge.
● Bunsen burner.
● Density meter, Ultrospec^TM^ 10 Classic (Biochrom).
● Magnetic specimen holder (Okolab) and magnetic sample locks (Okolab).
● Pipettes (Gilson).
● Plasma cleaner, PDC-002-HP (Harrick Plasma).
● Shaker incubator (Innova 42).
● Sonicating water bath, J. P. Selecta^TM^ ultrasons-H (Fisher scientific).

### 2.2 Supplies

● 18 x 150 mm sterile culture tubes with lid.
● 90 mm Petri Dishes (Fisher scientific).
● Attofluor^TM^ chambers (Invitrogen).
● Coverslips thickness 1.5H (0.17 ±0.005 mm round, 25 mm) (Marienfeld) (see **Note 2**).
● Disposable 1 mL cuvettes (Fisher scientific).
● Immersion Oil type F2 (Nikon).
● Reverse osmosis-purified water (Millipore MilliQ water purification system).
● Stericup^®^ 500 mL vacuum filters (Merck).
● Sterile 50 mL conical tubes (Falcon) and 2 mL tubes (Eppendorf).
● TetraSpeck^TM^ Microspheres, 0.1 µm, fluorescent blue/green/orange/dark red (Invitrogen).

### 2.3 Reagents

● 2-Mercaptoethanol (Sigma-Aldrich).
● Agar (Becton Dickinson).
● Amino acids (Sigma-Aldrich).
● Bromophenol blue (Sigma-Aldrich).
● Concanavalin A (ConA) (Sigma-Aldrich).
● D (+)-Glucose (Sigma).
● Dimethyl sulfoxide (DMSO) (Sigma-Aldrich).
● Dithiothreitol (Sigma-Aldrich).
● D-Sorbitol (VWR).
● Glycerol (Fisher Scientific).
● Latrunculin A (LatA) (Enzo Life Sciences).
● Magnesium chloride (MgCl_2_) (Fisher Scientific).
● Protease inhibitor cocktail (Sigma-Aldrich).
● Rapamycin (Rap) (Sigma-Aldrich).
● Sodium azide (NaN_3_) (Sigma-Aldrich).
● Sodium chloride (NaCl) (Merck).
● Sodium Dodecyl Phosphate (SDS) (Sigma-Aldrich).
● Sodium fluoride (NaF) (Sigma-Aldrich).
● Synthetic complete mixture drop-out without tryptophan, leucine, histidine and uracil (Formedium).
● Tris Base (Fisher Scientific).
● Triton X-100 (Sigma-Aldrich).
● Yeast Nitrogen Base (YNB) without amino acids (Formedium).
● YNB - Low Fluorescence, without amino acids, folic acid, and riboflavin (LoFlo) (Formedium).
● YNB without amino acids and ammonium sulphate (Formedium).
● YPD (Formedium).
● Zymolyase (Zymo Research).

### 2.4 Yeast strains

● Y7039 (*MATα, his3Δ1 leu2Δ0 ura3Δ0 LYS2+, can1Δ::Ste2pr-Leu2, lyp1Δ::*) (**32**).
● BY4741 (*MATa, his3Δ1, leu2Δ0, met15Δ0, ura3Δ0*) (Invitrogen).
● MKY2128 (*MATα, his3Δ1, leu2Δ0, ura3Δ0, LYS2+, can1Δ::STE2pr-LEU2, Lyp1Δ::, tor1-1, fpr1Δ::klUra*) (**7**).
● Y05143 strain (*MATa, his3Δ1 leu2Δ0 ura3Δ0, met15Δ0, sec6-4::kanMX4*) (**33**).

### 2.5 Primers for *tor1* rapamycin-resistance point mutation

The mutated nucleic acid is indicated in lowercase and bold.

**Forward primer**:

ATCAGAGTAGCCGTTCTATGGCACGAATTATGGTATGAAGGACTGGAAGATGC

GAG**a**CGCCAATTTTTCGTTGAACATAACATAGAAAAAATGTTTTCTACTTTAG

AACC

**Reverse primer**:

GGTTCTAAAGTAGAAAACATTTTTTCTATGTTATGTTCAACGAAAAATTGGCG**t**

CTCGCATCTTCCAGTCCTTCATACCATAATTCGTGCCATAGAACGGCTACTCTG

AT

## 3. Methods

The time estimates for each step are enclosed within brackets.

### 3.1 Culture media preparation (∼ 2 days)

1. YPD Medium (1 % yeast extract, 2 % peptone, 2 % glucose): dissolve 50 g of YPD in MilliQ water and adjust the volume to 1 L. For solid YPD medium, add 20 g of agar to the above solution. Autoclave and store the liquid medium at room temperature (RT), and the solid medium, once poured on sterile petri dishes, at 4 °C (see **Note 3**).
2. D (+)-Glucose 20 % w/v Solution: Dissolve 200 g of D (+)-Glucose in MilliQ water and adjust the volume to 1 L. Sterilise the solution through filtration using Stericup® or a 0.22 µm sterile membrane, and store at RT.
3. LoFlo Medium: Weigh 6.9 g of YNB without amino acids, folic acid, and riboflavin, and 1.9 g of amino acid DO Trp-. Add 900 mL of MilliQ and autoclave. Once the solution has cooled, add 100 mL of sterile D (+)-Glucose 20 %, mix with a sterile stirrer, and store at RT.
4. Synthetic Defined (SD) medium: Weigh 6.9 g of YNB without amino acids, 20 g of agar, 1.4 g of synthetic complete mixture drop-out and 0.1 g of desired additional amino acids (see **Note 4**). Add 900 mL of MilliQ water and autoclave. Add 100 mL of sterile D (+)-Glucose 20 %, mix with a sterile stirrer, pour in sterile petri dishes and store at 4 °C (see **Note 3**).

### 3.2 Construction of PICT strains (∼ 2 months)

The user can obtain the strains described in this step upon request from the authors.

1. The *S. cerevisiae* PICT strains derive from the Y7039, which is engineered to perform Synthetic Genetic Array (SGA) applications (**32**), and the BY4741 (Invitrogen). The genotype of these strains is detailed in step 2.4.
2. To obtain the *tor1-1* rapamycin resistant strains insert the S1972R point mutation into the *TOR1* gene by homologous recombination (**34**) using the primers listed in step 2.5. Select positive colonies on YPD plates containing 100 nM rapamycin and confirm by sequencing (*see* **Note 5**).
3. To delete the endogenous *FPR1* gene, replace it with the *klURA* cassette by homologous recombination (**34**). Select positive colonies on SD Ura- plates and confirm by sequencing (*see* **Note 5**).
4. To construct the FKBP/FRB system use homologous recombination on the Y7039 *tor1-1* and *fpr1Δ* derivative strain to sequentially fuse the RFP-FKBP tag to the Sla2 C-terminus and the FRB tag to the Exo70 C-terminus (**7**). Confirm the correct chromosomal integration by PCR.
5. To construct the prey-mNG strains use homologous recombination on BY4741 harbouring the *tor1-1* mutation to fuse the mNG tag to the Sec5 and the Sec2 C-terminus. Confirm the correct chromosomal integration by PCR.
6. Finally, use the SGA methodology to mate the Y7039 derivative strain from step 3.2.4 (*MATα, can1Δ::STE2pr-LEU2, Lyp1Δ::, his3Δ1, leu2Δ0, ura3Δ0, LYS2+, tor1-1, fpr1Δ::klUra, SLA2-RFP-FKBP::natNT2, EXO70-FRB::hphNT1*) with the corresponding BY4741 derivative strain from step 3.2.5 (*MATa, tor1-1, prey-mNG::kanMX4*). Select the haploid strains expressing Sla2-RFP-FKBP, Exo70-FRB and the desired prey-mNG. A detailed protocol for this approach can be found in references **32** and **35** (see **Note 6**). As a proof-of-concept we generated two strains:

● OGY1322 (*MATa, can1Δ::STE2pr-LEU2, Lyp1Δ::, his3Δ1, leu2Δ0, ura3Δ0, LYS2+, tor1-1, fpr1Δ::klUra, SLA2-RFP-FKBP::natNT2, EXO70-FRB::hphNT1, SEC5-mNG::kanMX4)*
● OGY1323 (*MATa, can1Δ::STE2pr-LEU2, Lyp1Δ::, his3Δ1, leu2Δ0, ura3Δ0, LYS2+, tor1-1, fpr1Δ::klUra, SLA2-RFP-FKBP::natNT2, EXO70-FRB::hphNT1, SEC2-mNG::kanMX4*).

### 3.3 Functional validation of the PICT strains

To rule out possible artefacts derived from genetic engineering it is necessary to evaluate cell growth and secretion in the PICT strains. Measuring the doubling time is a common approach to assess cell fitness in yeast (**36**). The endo-β-1,3-glucanase (Bgl2) secretion assay provides a specific readout to evaluate the functional aspect of exocytosis (**37**).

#### 3.3.1 Doubling time assay (∼ 2 days)

Day 1

1. Inoculate the PICT yeast strains and the growth control strain MKY2128 into 5 mL of YPD medium. Incubate the cultures in an orbital shaker at 200 rpm overnight (ON) at 25°C.

Day 2

2. Determine the cell concentration by measuring the optical density of the culture at 600 nm (OD_600_) in a spectrophotometer. Dilute the cultures to an OD_600_ of 0.1 in 40 mL of YPD. Incubate the cultures in an orbital shaker at 200 rpm at 25 °C.

3. After 3 hours, measure the OD_600_ of the culture again.

4. Repeat the previous step 3.3.1.3 at regular intervals of 1.5 hours.

5. Plot the OD_600_ values against time and fit a growth curve. Determine the doubling time of the yeast culture. The doubling time of the PICT strains should not be significantly different from the control strain (**Fig. 1**).

**Fig. 1.**
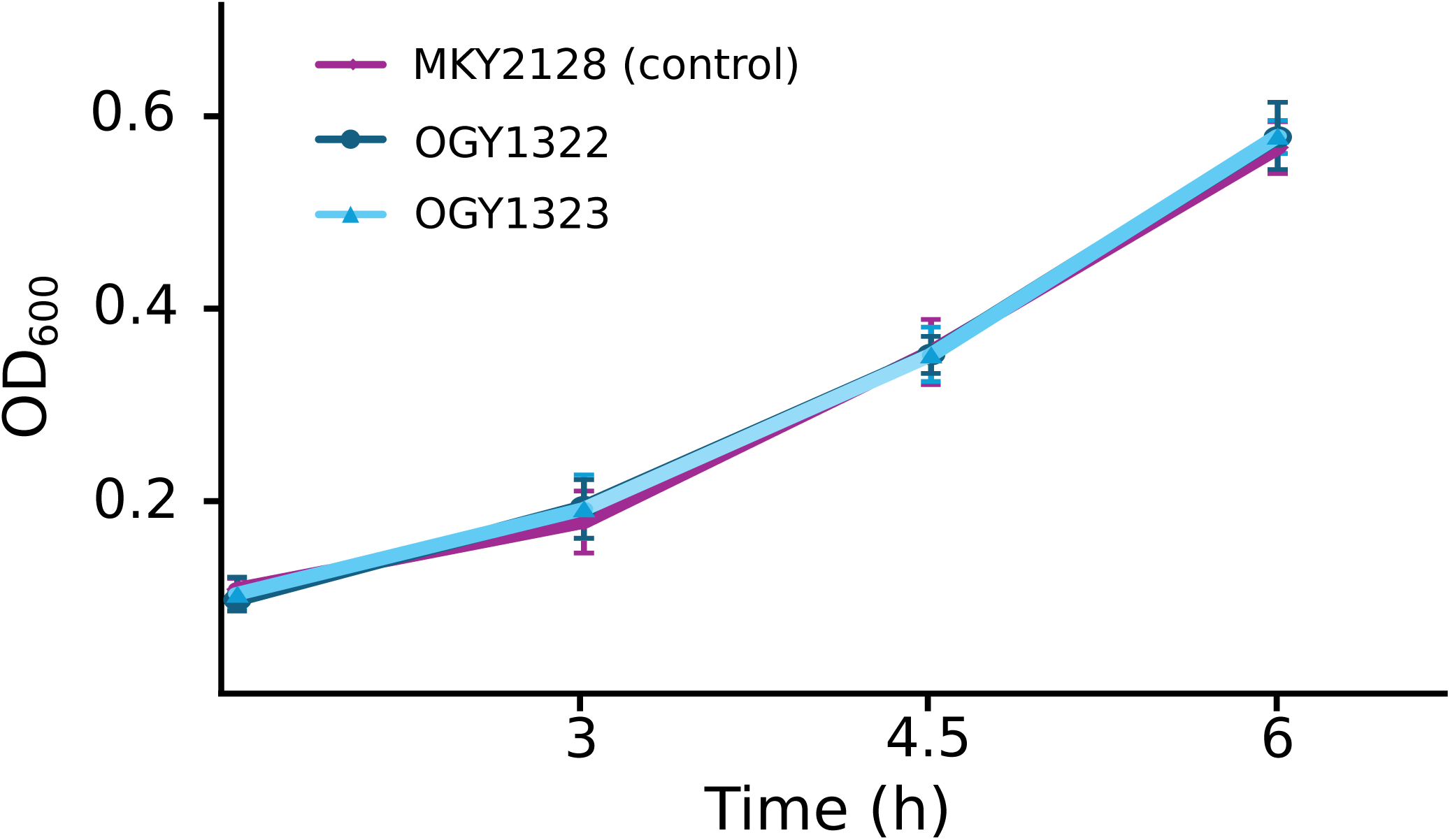
Doubling time assay. Doubling time is calculated using 3 and 6 hours’ time points from 3 biological replicates. No significant differences are observed between the mean doubling times of strains MKY2129 (control), OGY1322, and OGY1323 (1.8 ± 0.2, 1.9 ± 0.2, and 1.9 ± 0.2 hours, respectively). The difference between mean doubling times is assessed with a student’s t-test (p value > 0.05).

### 3.3.2 Bgl2 secretion assay (∼ 2 days)

This protocol follows the approach outlined in a recent study (**38**) with minor modifications.

Day 1

1. Inoculate the PICT yeast strains and the secretion control strain Y05143 into 5 mL of YPD medium (grow two separate cultures of the Y05143 strain). Incubate the cultures in an orbital shaker at 200 rpm at 25°C ON.

Day 2

2. Dilute the cultures to an OD_600_ of 0.1 in 20 mL of YPD. Incubate in an orbital shaker at 200 rpm at 25 °C for ∼ 5-6 hours (until OD_600_ 0.6–0.8). Shift one of the Y05143 strain cultures to 37°C for additional 2 hours (see **Note 7**).

3. Add NaN_3_ and NaF directly to the cultures at a final concentration of 10 mM each one. Transfer to a 50 mL conical tube and spin down the cells by centrifugation at 2,500 rpm at RT for 5 minutes.

4. Discard the remaining supernatant and resuspend the pellet in 1 mL of wash buffer (20 mM Tris–HCl, pH 7.5, 10 mM NaN_3_, and 10 mM NaF), transfer to a 2 mL Eppendorf tube. Wash the sample twice with 1 mL of wash buffer, followed by centrifugation at 2500 rpm at RT for 5 minutes.

5. Discard the remaining supernatant, resuspend the pellet in 300 µL of spheroplast solution buffer (50 mM Tris–HCl, pH 7.5, 1.4 M sorbitol, 10 mM NaN_3_, 10 mM NaF, 30 mM 2-Mercaptoethanol, and 0.2 mg/mL zymolyase) and incubate at 37 °C in a water bath for 30 minutes (during this incubation the cell wall is hydrolysed to form spheroplasts).

6. Gently spin down the spheroplasts by centrifugation at 1000 rpm at 4 °C for 5 minutes. Carefully transfer 100 µL of the supernatant to a new tube and mix with 20 µL of 6x SDS sample buffer (300 mM Tris–HCl, pH 6.8, 600 mM dithiothreitol, 12 % sodium dodecyl sulphate, 0.6 % bromophenol blue, and 60 % glycerol). This is the extracellular pool.

7. Gently aspirate the remaining supernatant using a pipette and wash the spheroplasts twice with 1 mL of spheroplast washing buffer (50 mM Tris–HCl, pH 7.5, 1.4 M sorbitol, 10 mM NaN_3_, and 10 mM NaF) to remove residual extracellular pool.

8. Lysate the spheroplasts by adding 300 µL of lysis buffer (20 mM Tris–HCl, pH 7.5, 100 mM NaCl, 2 mM MgCl_2_, 0.5 % Triton X-100, and 1x protease inhibitor cocktail) and incubate in ice for 10 minutes.

9. Remove the cell debris by centrifugation at 13,000 rpm at 4 °C for 5 minutes. Transfer 100 µL of the supernatant to a new tube and mix with 20 µL of 6x SDS sample buffer. This is the intracellular pool.

10. Heat the intracellular and extracellular pool samples for 5 minutes at 95 °C. Centrifuge very briefly in a microcentrifuge, load 5 µL of each sample and run in a 12 % SDS-PAGE gel.

11. Detect Bgl2 in a western blot using an anti-Bgl2 antibody (our antibody was a gift from Prof. Randy Schekman), diluted at 1:2,000 in TBST (1x Tris-Buffered Saline, 0.1% Tween^®^ 20 Detergent) with 5 % milk (**Fig. 2A**). Quantify band intensities using densitometry with ImageJ Gel Analyzer and normalise Bgl2 values against input values (Ponceau S. staining) (**Fig. 2B)**.

**Fig. 2.**
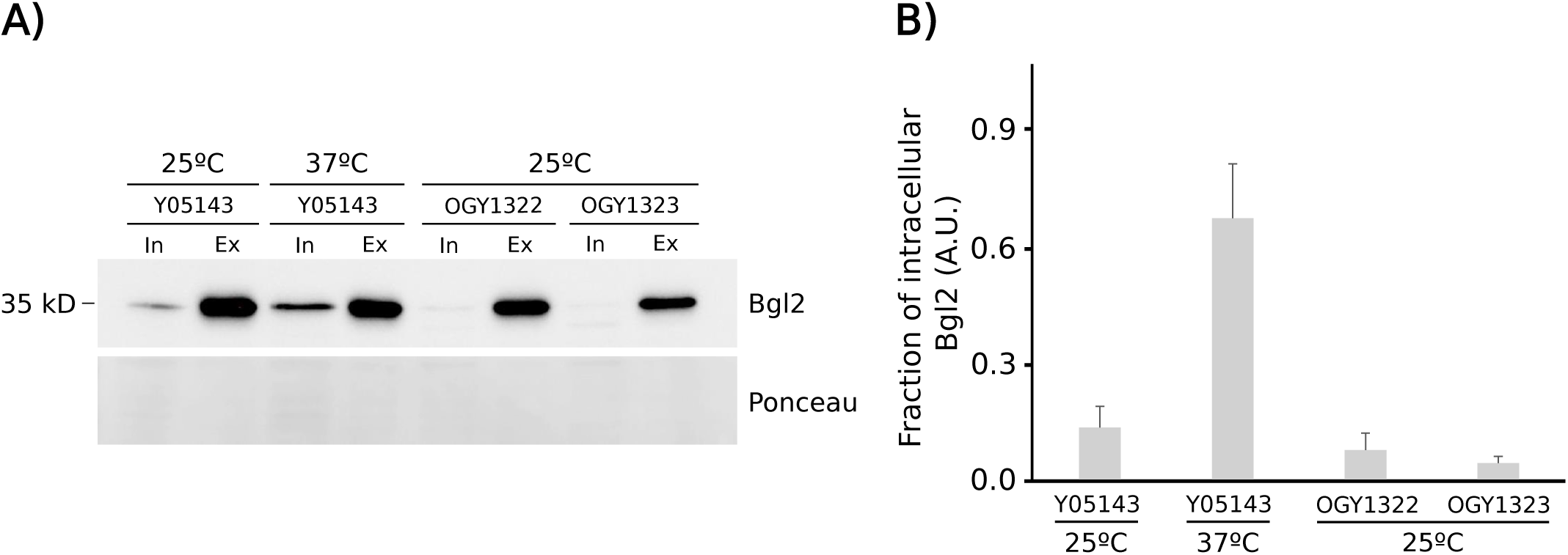
Bgl2 secretion assay. Mid-log phase growing Y05143 cells (control) are incubated for 2 hours at 25 °C or 37 °C. OGY1322 and OGY1323 strains are grown at 25 °C. **A**) Intracellular (In) and extracellular (Ex) pools of Bgl2 are quantified by western blot analysis (top). The total protein on the nitrocellulose membrane is stained with Ponceau S. (bottom). **B**) The ratio between intracellular and extracellular Bgl2 (In/Ex) was calculated from respective western blot band intensity measurement in three biological replicates. Error bars represent the standard deviation (n = 3). A.U.: arbitrary units.

### 3.4 Sample preparation for PICT experiments (∼ 2 days)

Day 1

1. Inoculate the PICT strains into 5 mL of YPD liquid medium and incubate the culture at 200 rpm in an orbital shaker at 25 °C ON.

Day 2

2. Determine cell concentration and dilute the cultures to an OD_600_ of 0.1 in LoFlo. Incubate the diluted cells in an orbital shaker at 200 rpm at 25 °C for ∼ 5-6 hours. The cells are ready for imaging when OD_600_ is between 0.6 and 0.8 (*see* **Note 8**). * We recommend dedicating these 5 hours to follow the beads imaging step (3.5).

3. Coat the coverslip surface with 50 µL of ConA (1 mg/mL in PBS). After 20 minutes of incubation at RT, wash the coverslip three times with LoFlo.

4. Add 50 µL of yeast culture on the coated coverslip and let the cells settle down at RT for 20 minutes.

5. Wash the cells three times with LoFlo (*see* **Note 9**). Thereafter, cells are ready for treatment.

6. Remove the LoFlo and add 50 µL of LoFlo with 10 µM rapamycin (stock solution at 2.47 mM in DMSO). Incubate at RT for 10 minutes.

7. Remove the LoFlo with 10 µM rapamycin and add LoFlo with 10 µM rapamycin and 200 µM LatA (stock solution at 20 mM in DMSO). After 10 minutes incubation cells are ready for imaging (*see* **Note 10**).

8. Put the coverslip inside the Attofluor^TM^ chamber and place it on the magnetic specimen holder. Secure it with the magnetic sample lock to minimise drift. Acquire the images following the instructions provided in step 3.6.

### 3.5 Beads imaging (∼ 30 minutes)

In all optical systems, chromatic aberration (CA) arises from variations in refractive index among optical components, causing the light wavelengths to focus at slightly different angles (**39**). Multicolour fluorescence imaging requires the correction of CA to prevent potential drawbacks such as reduced image resolution and inaccurate representation of the spatial distribution of fluorescent labels (**40**). Achieving precise CA correction involves the use of fluorescent beads to digitally re-align different wavelength channels with sub-pixel accuracy.

1. Use compressed air to eliminate any dust particles present on the coverslips where TetraSpeck™ Microspheres (hereafter beads) will be imaged. Subsequently, proceed to clean the coverslip using a plasma cleaner at 45 W and 200–500 mTorr for 1 minute (*see* **Note 11**).

2. Prepare a 1/100 dilution of beads in a 1:1 water/ethanol solution in an Eppendorf tube.

3. Thoroughly mix the beads by vortexing for 1 minute. Then sonicate for 15 minutes and finally vortex again for 1 minute (*see* **Note 12**).

4. Dispense 10 µL of the beads’ suspension onto the centre of a plasma-cleaned coverslip.

5. Allow the coverslip to air dry completely.

6. Position the coverslip within the Attofluor^TM^ chamber and add 50 µL of LoFlo (*see* **Note 13**). Position the Attofluor^TM^ chamber on the magnetic specimen holder and secure it with magnetic sample locks.

7. Perform the imaging on the focal plane containing the glass-attached beads using the red or green fluorescence channel (hereafter channel 1 and channel 2 respectively) and restrict the region of interest (ROI) to a homogeneously illuminated area of the camera sensor. Illumination intensity may vary substantially from the centre of the image to the periphery, therefore for our microscopy setup, the ROI is restricted to an area of 1024 x 1024 pixels (*see* **Note 14**). Use µManager to select the ROI either manually (drawing a 1024 x 1024 pixels box) or use the following path: “Menu” > “Edit” > “Selection” > “Specify” (With: 1024, Height: 1024, X and Y coordinates depends on the light distribution on the camera sensor). It is essential to optimise the light power to prevent saturation while maximising the photons detected in each channel (*see* **Note 15**).

8. Find a ROI with homogeneously distributed beads (∼ 25 - 75 beads in a 1024 x 1024 pixels ROI) (**5**). Then create a 10 x 10 grid in µManager by following this path: “Tools” > “Stage Position List” > “Create Grid” > “Tile creator” (“Overlap”: 90 %, “Pixel Size (µm)”: 0,065, create a 10 x 10 grid size and press the “Center Here” button), which generates a “Stage Position List” with 100 positions.

9. Configure the microscope to introduce a delay after each stage movement. This delay allows the system to stabilise before capturing the images, minimising any potential drift. In our system, we use a custom script by following this path: “Tools”

> “Script Panel” > “Run”.

import ij.gui.GenericDialog;

mm.acquisitions().clearRunnables();

*runnable = new Runnable() {*

int count = 0;

public void run() {

mmc.sleep(2500);

mm.scripter().message(“Waited 2.5 secs, nr: ” + count);

count++;

}

};

mm.acquisitions().attachRunnable(0, -1, 0, 0, runnable);

10. Acquire images. In our microscopy setup we use an exposure time of 2,500 ms and 20 % of LED power for each channel. The camera operates at 16 bits and 200 MHz pixel readout (lowest noise), with no binning.

### 3.6 Cells imaging (∼ 1 hour for the first strain and 10 additional minutes for each subsequent strain)

After beads imaging, acquire cell images using identical settings, except for the LED power which will be adjusted to prevent camera sensor saturation while maximising the detected photons in each channel. To minimise CA, do not change the filter cube during the imaging acquisition, including the acquisition of the corresponding brightfield (transmitted light) image (*see* **Note 16**).

1. Use the brightfield to locate a region of the sample where most of the cells are focused on their equatorial plane (with our ROI of 1024 x 1024 pixels, the expected number of cells is ∼ 15 cells/image). We image each channel sequentially (channel 1, RFP – channel 2, mNG). With an exposure time of 2,500 ms and 100 % LED power (*see* **Note 17**).
2. Move the stage as much as necessary to avoid imaging cells that had been illuminated during the acquisition of previous images (to avoid photobleaching). Capture as many images as possible within the 10-minutes window following the LatA treatment (refer to step 3.4.7). Collect the necessary amount of images to obtain around 300 cells with colocalising spots (e.g, 20 images for ∼ 15 cells / image). If necessary, prepare more than one sample for each bait-FRB - prey-mNG combination. **Fig. 3** illustrates an example of cells imaged before and after treatment with rapamycin and LatA.

**Fig. 3.**
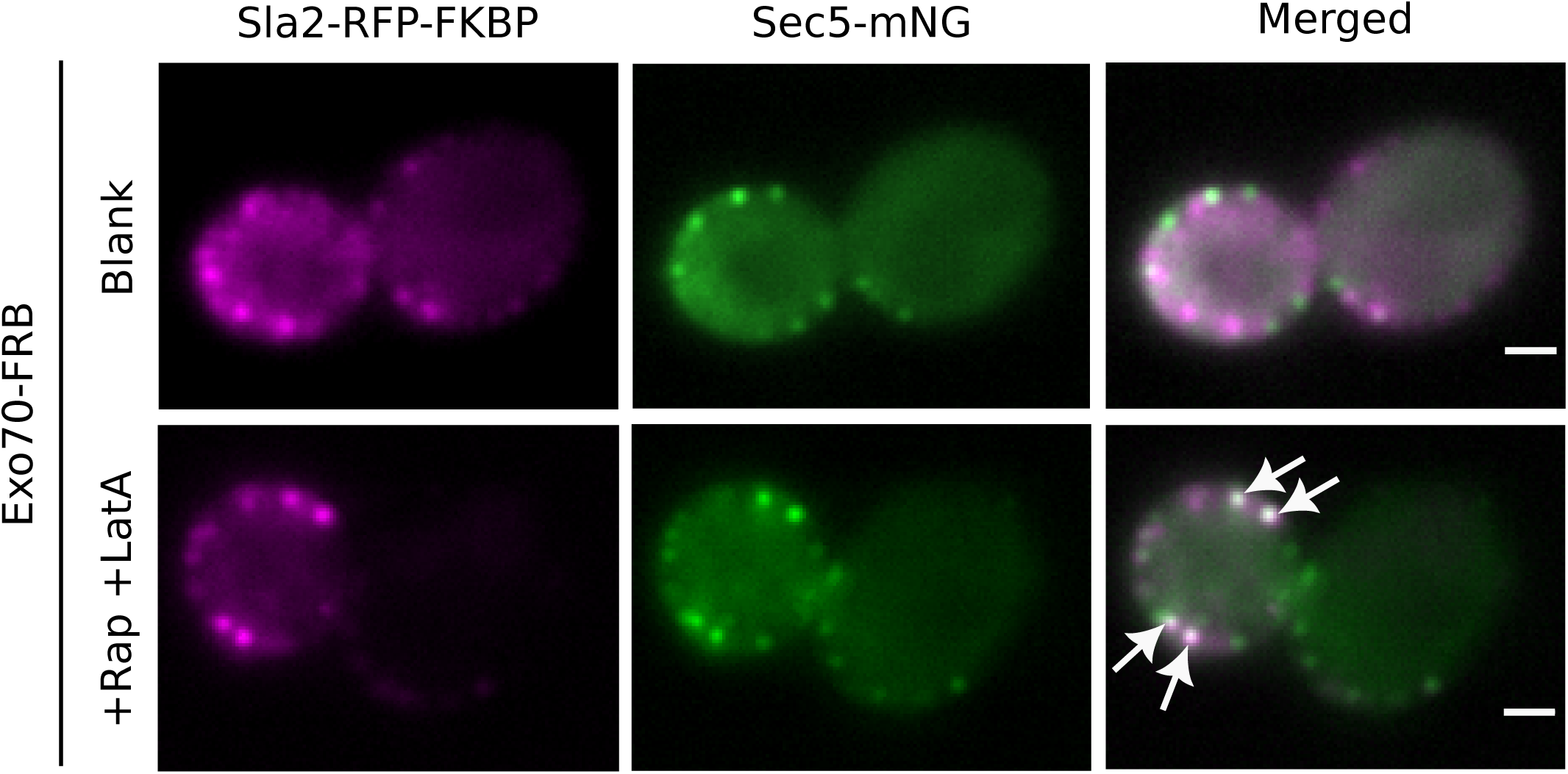
PICT experiment for the exocyst. Representative images of OGY1322 yeast cells expressing Sla2-RFP-FKBP anchoring platforms (left column, magenta) and Sec5-mNG prey protein (centre column, green) before (top) and after (bottom) treatment with rapamycin (Rap) and latrunculin A (LatA) using Exo70-FRB as bait. Arrows point at Sec5-mNG spots that co-localise with Sla2-RFP-FKBP. Scale bars = 1 µm.

### 3.7 Image analysis

Image analysis involves five key steps: 1) calculation of the image registration map, 2) image pre-processing, 3) spot detection, spot-pair linking and CA correction, 4) spot-pair filtering, and 5) outlier rejection and distance estimation. To simplify the image analysis protocol, we developed the PyF2F software (**41**), accessible at https://github.com/GallegoLab/PyF2F.git. PyF2F can run standalone on a Linux/Mac laptop from the command line or online using a Colab notebook (https://colab.research.google.com/drive/1kSOnZdwRb4xuznyQIpRNWUBBFKms91M8?usp=sharing) (recommended for Windows users).

#### PyF2F Standalone (Linux/MacOS) (∼ 18 minutes)

Below are the instructions for installing the PyF2F software and the steps for processing and analysing the PICT images via the command line. We demonstrate the step-by-step protocol by estimating the anchor-to-prey distance in images captured for the strains OGY1322 and OGY1323 (refer to step 3.2.6). We provide the recommended parameters to run PyF2F in Linux/Mac terminal given the optical system described above (refer to step 2.1). We suggest using a Conda or Miniconda package management system (*Anaconda Software Distribution*. Computer software. Vers. 2-2.4.0. Anaconda, Nov. 2016. Web.) to efficiently set up the PyF2F requirements.

The lines beginning with “>” represent commands to be executed on a UNIX-style command prompt.

#### 3.7.1 PyF2F installation (∼ 5 minutes)

1. Download the GitHub repository (*see* **Note 18**): > git clone https://github.com/GallegoLab/PyF2F.git > cd PyF2F
2. Download the CNN weights from https://zenodo.org/records/3598690. Alternatively, use the following commands (*see* **Note 19**): > pip install zenodo-get > zenodo_get -r 3598690
3. Create a Conda environment with Python version 3.7 named “pyf2f”: > conda create -n pyf2f python=3.7 anaconda > conda activate
4. Install the requirements listed in the *requirements.txt* file located in the /PyF2F/ directory: > pip install -r requirements.txt
5. Activate the pyf2f environment and change directory to the /scripts/ directory: > conda activate pyf2f > cd scripts/
6. Run a short test to quickly check that PyF2F runs without problems and the expected/output/ folder is generated (*see* **Note 20**). > bash run_pyf2f.sh ../short_example/ all

#### 3.7.2 Calculation of the image registration map (∼ 3 minutes)

As explained in step 3.5, CA between the two-channel images can be corrected using an image registration approach that requires imaging fluorescent beads (*see* **Note 21**). Initially, PyF2F calculates the registration map by applying an affine transformation to the images of beads located in /input/reg/ (**Fig. 4**). Subsequently, PyF2F assesses the target registration error (TRE) or registration accuracy using the images of beads located in /input/test/ (**Fig. 5**). In your terminal, execute the following command (*see* **Note 22**):

**Fig. 4.**
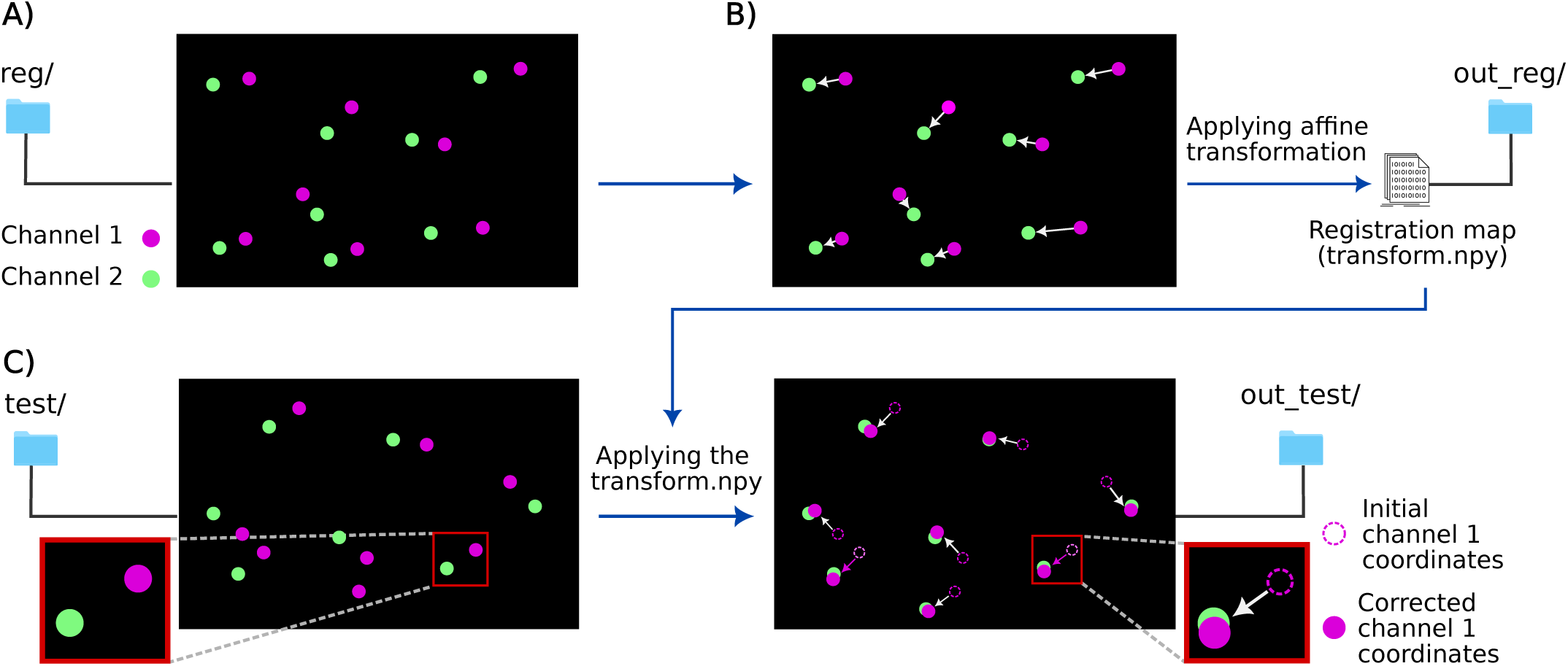
Illustration of the scheme to calculate the image registration map. **A)** A first set of images of beads (located in /reg*/)* is used to calculate the CA between the channel 1 (magenta dots) and the channel 2 (green dots). **B)** The registration map required to correct the CA between the two channels is calculated by applying an affine transformation to align the beads’ centroid coordinates of channel 1 towards the beads’ centroid coordinates of channel 2. The calculated registration map (“transform.npy”) is saved as a matrix in the directory /out_reg*/*. **C)** The registration map calculated in B) is employed to correct the beads’ centroid coordinates of channel 1 in a second set of bead images (located in /test*/*). This will be subsequently used to evaluate the registration accuracy (TRE, see Fig. 5).

**Fig. 5.**
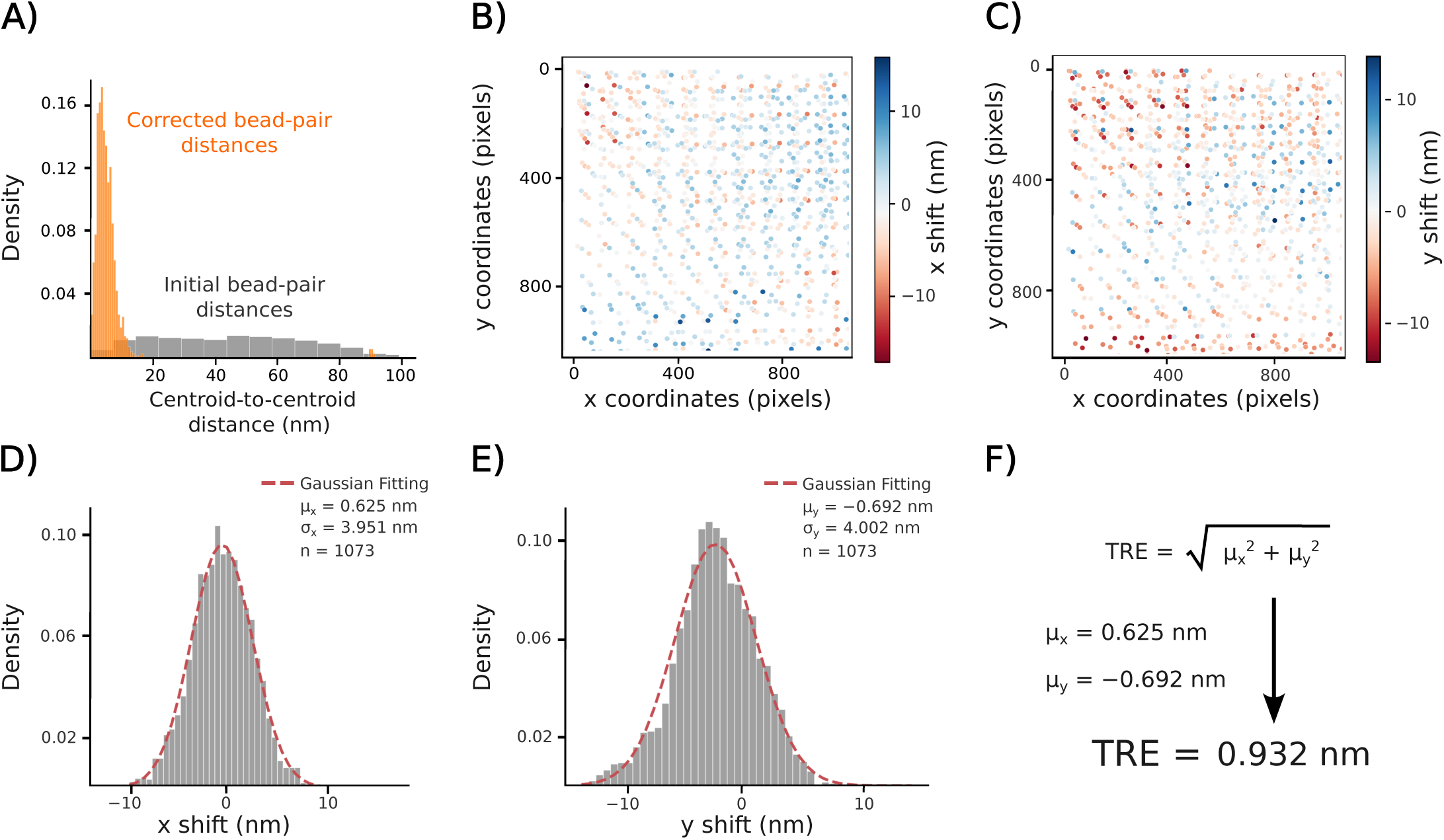
Calculation of the registration accuracy. **A)** The distribution of centroid-to-centroid distances between the two-channel beads’ centroid coordinates is represented before (grey) and after (orange) CA correction. **B, C)** Each dot represents a single bead, for which the difference (or shift) between the two-channel coordinates is calculated along the *x-*axis (**B**) and the *y*-axis (**C**) of a ROI image and is represented by the heat-map colours. **D, E**) The mean of the *x-*axis (µ_x_, **D**) and *y*-axis (µ_y_, **E**) deviations are the registration errors along each axis. **F)** The target registration error (TRE) can be calculated to estimate the accuracy of the registration. This registration was used for the subsequent analysis of Sec5-mNG and Sec2-mNG distances to Sla2-RFP-FKBP.

> bash run_image_registration.sh /working_directory/input/ reg test out_reg out_test

#### 3.7.3 Image pre-processing (∼ 3 minutes)

PyF2F corrects the background fluorescence of the PICT images (**Fig. 6A**) to precisely localise the centroid coordinates of diffraction-limited fluorescent spots in channel 1 and channel 2 (**Fig. 6B**) (*see* **Note 23**). In your terminal, execute the following command:

**Fig. 6.**
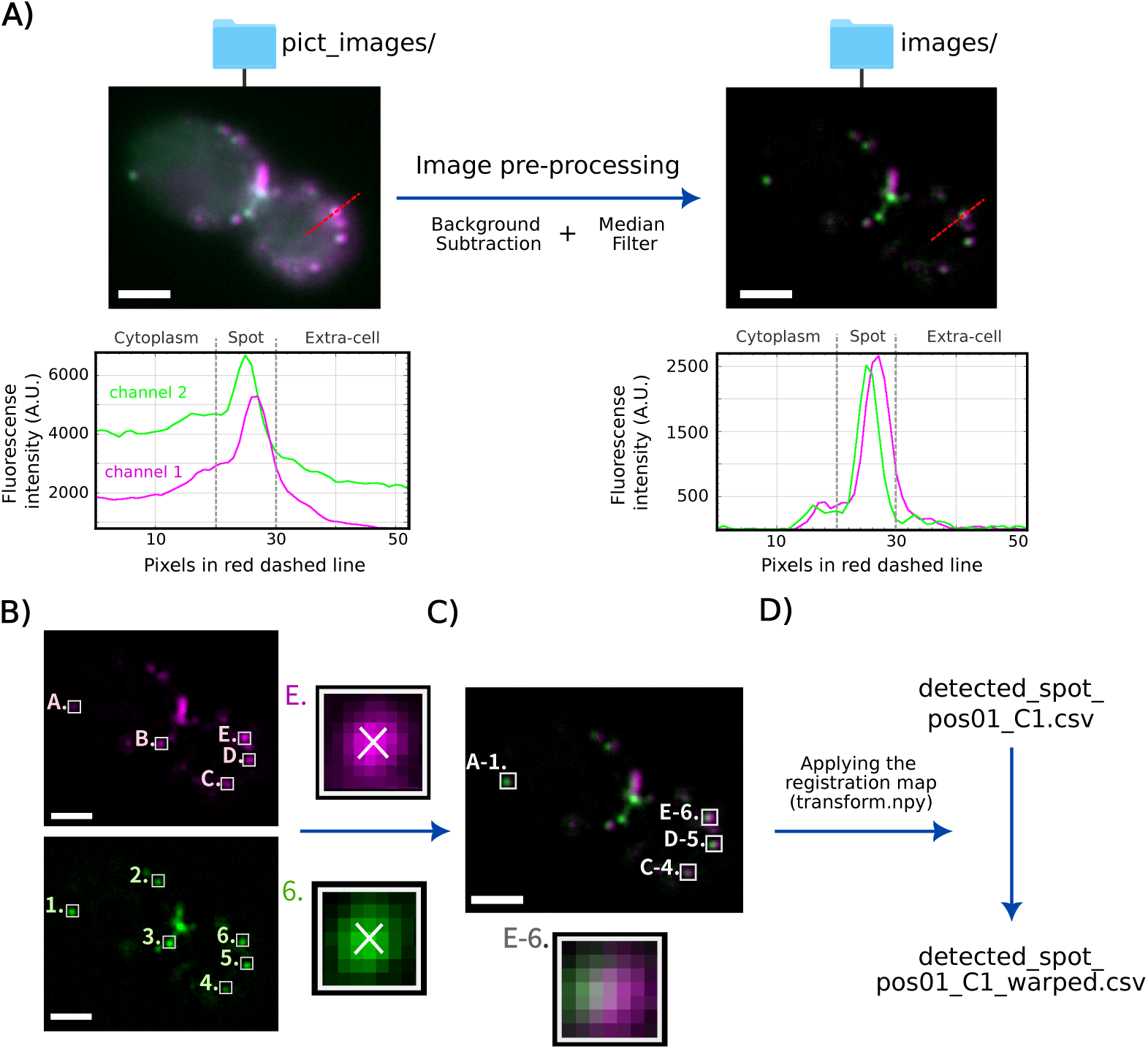
Illustration of the PICT image pre-processing, spot detection, spot-pair linking and CA correction. **A)** Raw two-channel PICT images located in /pict_images/ (top-left) are corrected for the extracellular (background subtraction) and intracellular (median filter subtraction) fluorescence and saved in /images/ (top-right). The figure shows an example of a *S. cerevisiae* cell (OGY1322 strain) before (left) and after (right) image pre-processing. The plots illustrate the fluorescence intensity profile along the red dashed line for channel 1 (magenta) and channel 2 (green) before (bottom-left) and after (bottom-right) image pre-processing. **B)** Diffraction-limited fluorescent spots are detected in channel 1 (i.e. spots A–E, magenta, top) and channel 2 (i.e. spots 1–6, green, bottom). The zooms-in show the centroid coordinates represented as a white cross (channel 1–spot E, top; channel 2–spot 6, bottom). **C)** Channel 1–channel 2 spot-pairs within a maximum separation set by the user are linked into “spot-pairs” (white boxes). The zoom-in shows an example of the spot-pair E–6 (bottom). **D)** The channel 1 centroid coordinates (in file detected_spot_pos01_C1.csv) are corrected for the CA using the registration map calculated in step 3.7.2 (transform.npy). Corrected coordinates are saved in a separate file (detected_spot_pos01_C1_warped.csv). Scale bars = 2 µm.

> bash run_pyf2f.sh /working_directory/ pp

#### 3.7.4 Spot detection, spot-pair linking and CA correction (∼ 30 minutes)

Co-localising diffraction-limited spots – channel 1 / channel 2 spots whose 2D centroid coordinates are located close together within a maximum radius set by the user – are linked into “spot–pairs” (**Fig. 6C**) (*see* **Note 24**). Then, the CA is corrected by using the registration map calculated in step 3.7.2 (**Fig. 6D**) (*see* **Note 25**). In your terminal, execute the following command:

> bash run_pyf2f.sh /working_directory/ spt,warping

#### 3.7.5 Spot-pair filtering (∼ 3 minutes)

Detected spot-pairs are sorted through the following consecutive steps: 1) selection of close-to-cell contour spot-pairs, 2) selection of high-quality spot-pairs and 3) selection of in-focus spot-pairs (**Fig. 7A-C**) (*see* **Note 26**). This filtering procedure is critical to reduce spurious spot-pairs before further analysis. In your terminal, execute the following command:

**Fig. 7.**
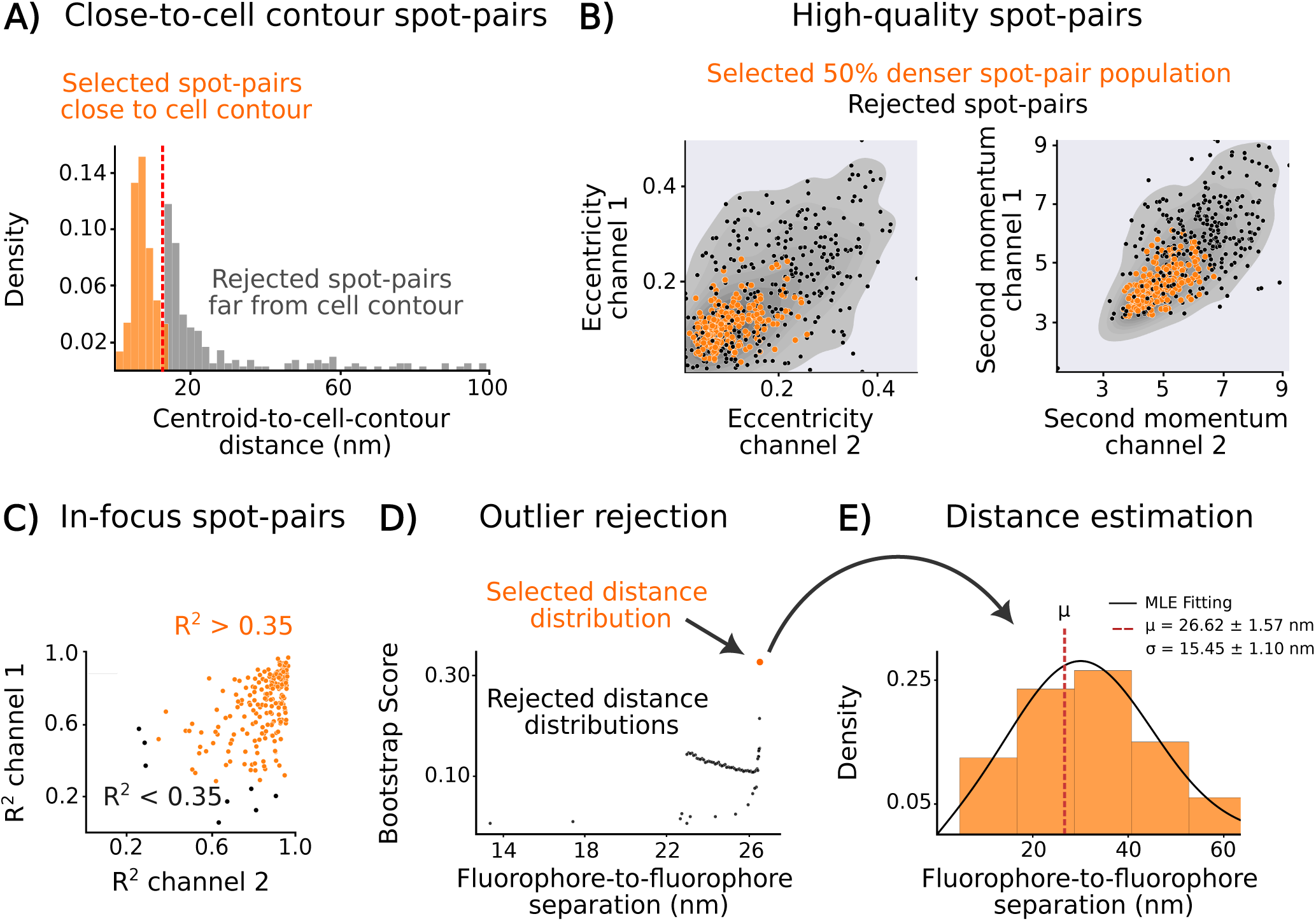
Filtering of spot-pairs, outlier rejection and estimation of the Sla2-RFP-FKBP to Sec5-mNG distance. Filtering of spot-pairs (**A-C**): **A)** Only spot-pairs close to the cell contour are selected (orange). The red-dashed line shows the cut-off to select or discard spot-pairs. **B)** Only spot-pairs in the 50% denser population based on intensity properties (eccentricity and second momentum) are selected (orange dots). **C)** Only in-focus spot-pairs (which intensity profile fits a 2D-Gaussian function) are selected (orange dots). Outlier rejection: **D)** After filtering, outliers are rejected using a bootstrap method (*see* **Note 27**). The black dots correspond to the bootstrap score of the iterative outlier rejections. The highest score (orange dot) corresponds to the distance distribution maximising the log-likelihood for µ and σ. Distance estimation: **E)** The figure shows the distance distribution corresponding to the best-scored distribution without outliers (n = 195) (orange dot in D). The MLE fitting gives a distance estimate µ = 26.62 ± 1.57 (σ = 15.45 ± 1.1) for the separation between the anchor Sla2-RFP-FKBP and the prey Sec5-mNG when the exocyst is anchored using Exo70-FRB as bait. Errors (±) represent the standard error (SE) of µ (SEµ) and σ (SEσ) estimations.

> bash run_pyf2f.sh /working_directory/ segment,kde,gaussian

#### 3.7.6 Outlier rejection and distance estimation (∼ 1 hours 30 minutes)

In this last step, PyF2F estimates the true average distance between a prey-mNG and Sla2-RFP-FKBP. Since the distribution of measured distances between paired diffraction-limited fluorescent spots follows a Rician distribution (**3, 4**), the true separation (µ) and standard deviation (σ) can be computed with a Maximum Likelihood Estimation (MLE) (**Fig. 7D**) (*see* **Note 27**). In your terminal, execute the following command:

> bash run_pyf2f.sh /working_directory/ mle

The bootstrap method aims to find the distance distribution that maximises the log-likelihood for estimating µ and σ given a Rice distribution (**3, 4**). This can be achieved by iteratively rejecting the data points that are less likely to belong to the dataset (false positives - outliers) and consequently minimise the log-likelihood. As shown in figure 7D and figure 7E, the bootstrap method scores the estimation for the average fluorophore-to-fluorophore distance (µ) after an outlier rejection (black dots), giving us a notion about the likelihood to find outliers in that distribution. The lower the score, the more likely to find outliers in that distribution. The highest scored estimate for the distance and standard deviation between Sla2-RFP-FKBP and Sec5-mNG centroid coordinates (using Exo70-FRB as bait) is µ = 26.62 ± 1.57 nm and σ = 15.45 ± 1.1 nm (errors (±) represent the standard error (SE) of µ and σ estimations) (**Fig. 7E**).

#### 3.7.7 Inter-complex distances with PICT

Besides measuring distances within a protein complex, PICT also facilitates the quantitative assessment of transient inter-complex interactions. This capability allows us to investigate the spatial organisation of the exocyst and its transient protein-protein interactions. To illustrate this approach, we investigated the spatial organisation of the exocyst and Sec2, a guanyl-nucleotide exchange factor that associates with the surface of secretory vesicles (**Fig. 8**). To be functional, the exocyst complex requires the binding of the Sec15 subunit to Sec2 (**12, 26, 27**). The spatial distribution of the 23 ± 9 copies of Sec2 that, on average, populate each secretory vesicle remains unknown (**42**).

**Fig. 8.**
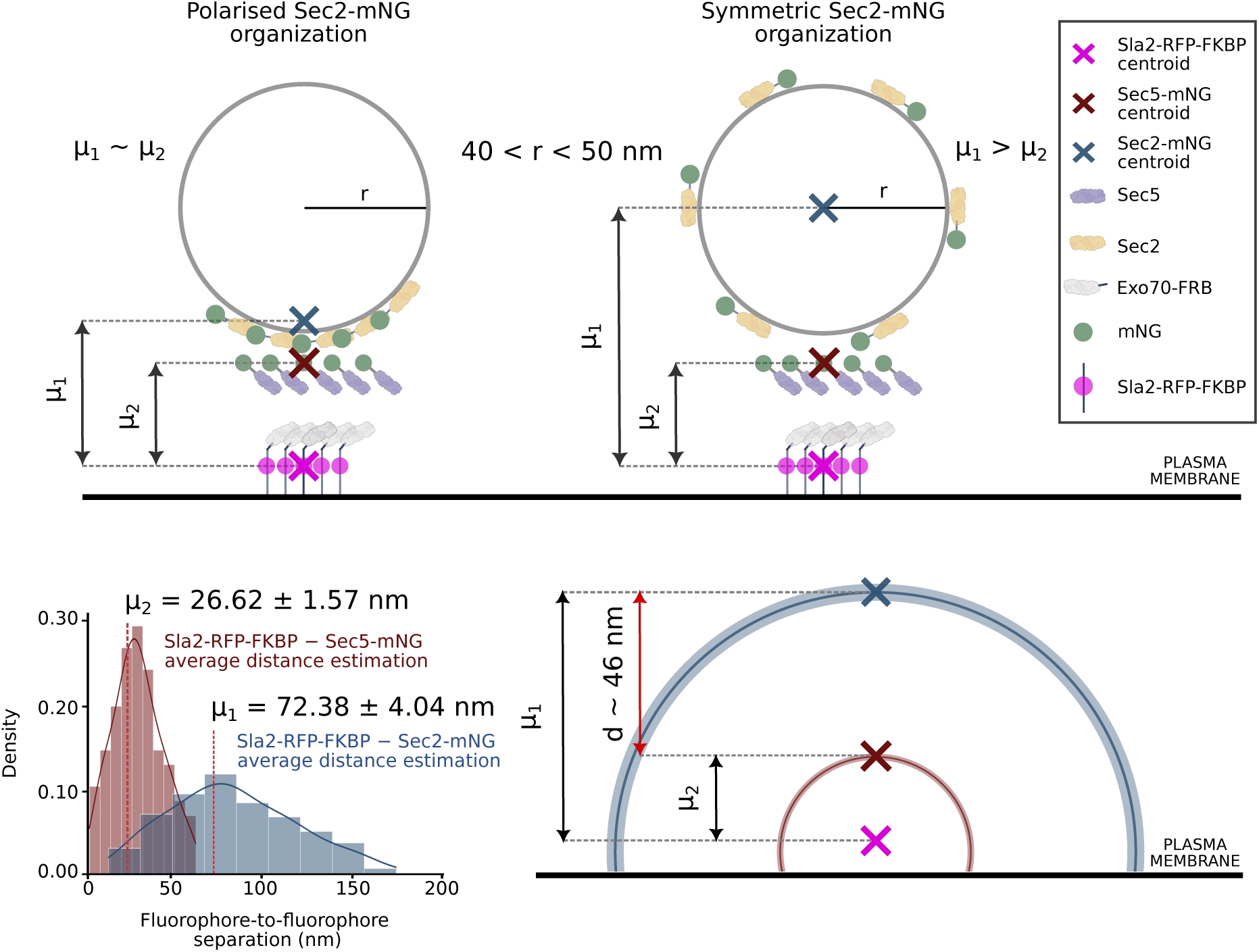
Distance estimation and spatial organisation of exocyst interactions. Top-left: Illustration of a model where, upon tethering, multiple Sec2-mNG cluster on the vesicle surface adjacent to the exocyst. In this model, the average separation between Sla2-RFP-FKBP and Sec2-mNG (µ_1_) is similar to the average separation between Sla2-RFP-FKBP and Sec5-mNG (µ_2_). Top-right: Illustration of a model where, upon tethering, multiple Sec2-mNG are symmetrically distributed throughout the vesicle surface. In this model the centroid coordinate of Sec2-mNG corresponds to the centre of the vesicle, about 40–50 nm from the surface where the exocyst is bound. Thus, the average separation between Sla2-RFP-FKBP and Sec2-mNG (µ_1_) is expected to be, at least, 40 nm longer than the average separation between Sla2-RFP-FKBP and Sec5-mNG (µ_2_). Bottom-left: Distance distribution for the separation between Sla2-RFP-FKBP and Sec2-mNG (µ_1_ = 72.38 ± 4.04; σ = 39.25 ± 2.91) and between Sla2-RFP-FKBP and Sec5-mNG (µ_2_ = 26.62 ± 1.57; σ = 15.45 ± 1.1). Errors (±) represent the standard error (SE) of µ (SEµ) and σ (SEσ) estimations. Bottom-right: Illustration of the spatial organisation derived from the PICT experiment: Sla2-RFP-FKBP platform associated to the plasma membrane acts as spatial reference; the exocyst subunit Exo70-FRB is anchored to Sla2-RFP-FKBP; the centroid coordinate of Sec2-mNG can be located at any point along a circumference of 72.38 ± 4.04 nm of radius (µ_1,_ blue line) from the Sla2-RFP-FKBP centroid coordinates; the centroid coordinates of Sec5-mNG can be located at any point along a circumference of 26.62 ± 1.57 nm of radius (µ_2_, green line) from the Sla2-RFP-FKBP centroid coordinates; the distance between the centroid coordinates of Sec5-mNG and the centroid coordinates of Sec2-mNG is about 46 nm (red arrow). This spatial organisation supports a model where Sec2-mNG remains symmetrically distributed on the surface of the tethered secretory vesicles.

In the present study we used the distance from the Sla2-RFP-FKBP spatial reference to Sec5-mNG and to Sec2-mNG to investigate the distribution of Sec2-mNG throughout the surface of the secretory vesicle, which has a diameter of 80 to100 nm (**14, 43**). The two measurements presented in this chapter define the average conformational space that Sec5-mNG and Sec2-mNG can occupy with respect to a spatial landmark (i.e., Sla2-RFP-FKBP where Exo70-FRB is anchored, illustrated in **Fig. 8, top**). If exocyst binding induces a major clustering of multiple Sec2-mNG towards the vesicle side adjacent to the tethering complex, the centroid of the Sec2-mNG diffraction-limited spot will be located near the vesicle-exocyst interface. Then, the average separation between Sla2-RFP-FKBP and Sec2-mNG (µ_1_) will be similar to the average separation between Sla2-RFP-FKBP and Sec5-mNG (µ_2_) (**Fig. 8, top-left**). Instead, if upon vesicle-exocyst binding the multiple Sec2-mNG remains symmetrically distributed around the vesicle surface, the average Sec2-mNG centroid coordinates will correspond to the vesicle centre (**Fig. 8**, **top-right**). In this case, the average Sec2-mNG centroid coordinates will be about 40–50 nm from the surface where the exocyst is bound, being µ_1_ > µ_2_.

We applied the protocol to measure the average separation between Sla2-RFP-FKBP and Sec5-mNG and between Sla2-RFP-FKBP and Sec2-mNG (always using Exo70-FRB as bait). The measured distances were 26.62 ± 1.57 nm and 72.38 ± 4.04 nm respectively (estimated distance (µ) ± standard error of µ (SEµ)) (**Fig. 8, bottom-left**). Therefore, Sec5-mNG is closer to the Exo70-FRB (both subunits are within the same protein complex) than Sec2-mNG. Remarkably, on average Sec2-mNG is ∼ 46 nm further away than Sec5-mNG (**Fig. 8, bottom-right**) supporting a model where Sec2-mNG remains symmetrically distributed on the surface of the tethered secretory vesicles with an average radius of 40–50 nm.

## 4. Notes

1. If the user employs fluorophores with substantial intensity spectrum overlap, adding a filter wheel could be beneficial to mitigate potential bleed-through. This adjustment is recommended rather than essential, according to the specific demands of your experiment.
2. Use of glass-bottomed dishes or rectangular coverslips is also possible.
3. When preparing solid media pour 20 mL of the autoclaved media into 90 mm sterile petri dishes. Lay the petri dishes on a flat surface before agar solidification. Allow them to dry for 1 day at RT to prevent condensation and store the plates upside down at 4 °C in a plastic bag.
4. If the strain’s selection medium includes a drug (e.g., for *kanMX4*, *hphNT1* and *natNT2* cassettes use the selection on Geneticin/G418, Hygromycin B and Nourseothricin/ clonNAT, respectively), use YNB without ammonium sulphate and add monosodium glutamate as the nitrogen source. After autoclaving, allow the medium to cool down to ∼ 60 °C before adding the antibiotic to prevent its degradation.
5. Rapamycin binds to endogenous Tor1 and Fpr1 proteins in wild-type yeast. To prevent undesired Tor1 inhibition, the rapamycin binding pockets in the FRB domain of Tor1 must be mutated (S1972R). To prevent undesired effects and competitive inhibition to rapamycin induced heterodimerization of FKBP and FRB tags, endogenous Fpr1 must be deleted.
6. SGA methodology consists of a series of replica pinning steps, which enable the construction of haploid double mutants through mating and meiotic recombination and can generate various mutant combinations in high throughput (**32**). We recommend its use due to the large number of strains required for various bait-prey pairings. Alternative yeast transformation could be used as well.
7. Y05143 is a strain that codes for *sec6-4*, a temperature-sensitive mutant of the exocyst subunit Sec6. Y05143 exhibits near-normal Bgl2 secretion at 25 °C but fails to perform exocytosis at the restrictive temperature of 37 °C.
8. It is important that, during growth, yeast cells stay in mid-log phase (OD_600_ = 0.6–0.8). At higher OD the carbon source might be depleted, which could lead yeast cells to undergo a transition to a growth mechanism based on aerobic respiration, known as the diauxic shift. During this process, yeast cells exhibit auto-fluorescence at different wavelengths, reduced expression of fluorescent proteins, and uneven fluorescence distributions (**44**).
9. Critical step: Withdraw superfluous medium and strongly rinse off nonadherent cells with LoFlo medium. It is very important to eliminate floating or incompletely adhered cells.
10. Critical step. To minimise artefacts derived from compromised cell fitness, do not extend the LatA treatment for more than 20 minutes before finishing image acquisition.
11. Cleaning the coverslips with plasma will facilitate a uniformly dispersed distribution of the fluorescent beads. If a plasma cleaner is not available, coverslips could be cleaned with ethanol or methanol. However, it’s important to note that when cleaning the glass with these solvents you may need to manually spread the droplet by pipetting, as the level of hydrophilicity achieved will be lower than using a plasma cleaner.
12. Beads tend to aggregate, which prevents optimal centroid localisation. Following the procedures is essential to minimise spurious CA correction.
13. To mimic the conditions under which the yeast cells are imaged, we image beads on the same coverslips, Attofluor™ chamber and LoFlo media.
14. Before imaging, make sure that the emission light from both channels is homogeneously detected in the selected ROI. If the uniformity of illumination is suboptimal, this will have a severe impact on the reliability of automated spot detection and centroid localization (**45**). For that purpose, you may restrict the image acquisition to the sensor area that minimises uneven detection of the fluorophores. To assess the evenness of light distribution across the field of view and its proper alignment, we utilise Chroma fluorescent plastic slides.
15. Due to the significantly higher fluorescence of beads compared to fluorescent proteins, it is essential to adjust the light power separately to not saturate the camera sensor. Achieving optimal bead imaging is crucial for generating an accurate CA correction (refer to step 3.7.2). Saturated signal will hamper centroid localization and, therefore, it will compromise centroid-to-centroid distance estimation.
16. When imaging multiple strains, it is practical that one person handles the sample preparation while another person images the cells. This division of tasks is crucial to preserve the time constraints following the LatA treatment (see step 3.4.7).
17. Settings of image acquisition depend on the imaging system, camera, acquisition software, and signal of the samples. Such long exposure maximises the number of photons collected while averaging a large conformational space of the anchored exocyst subunits. Using appropriate microscopy settings is important to obtain a good fluorescence image. In your system, adjust the light power to be optimal for the dynamic range of the camera used and the signal of the sample.
18. The git repository can also be downloaded as ZIP from the Gallegolab repository (https://github.com/GallegoLab/PyF2F).
19. PyF2F utilises the pre-trained weights for the convolutional neural network (CNN) that is used in Yeast Spotter for yeast cell segmentation (**46**). The weights are necessary to run the software, but are too large to share on GitHub. You can download the ZIP file from the following Zenodo repository (https://zenodo.org/record/3598690).
20. Notice that the PyF2F repository contains two examples to test the image analysis protocol. We recommend testing the PyF2F performance on the /short_test/ or /long_test/ cases before running PyF2F on your own data. The detailed guidelines for running PyF2F on both examples, as well as extended information about the content of the PyF2F repository, can be found at the Gallegolab repository (https://github.com/GallegoLab/PyF2F).
21. Notice the difference between the terms “image registration” and “CA correction”. “Image registration” is the process to geometrically align two or more images of the same scene (**47**). In this protocol, image registration is used to align the two-channel images of beads (using channel 2 as reference) to calculate the registration map. “CA correction” is the process to correct the spot centroid coordinates of channel 1 using the registration map previously calculated. In this protocol, CA correction is used 1) to calculate the registration accuracy using images of beads (step 3.7.2) and 2) to correct the spots’ centroid coordinates found in channel 1 of the PICT images (step 3.7.4).
22. PyF2F uses the bead images located in /reg/ to calculate the CA between the channel 1 and channel 2. In an ideal scenario (without CA), the beads’ *xy* coordinates should be exactly the same in each channel. However, CA deviates the beads’ coordinates of one colour channel with respect to the other. This optical aberration must be registered prior to calculating distances between the centroid of each colour signal. PyF2F calculates the registration map by applying an affine transformation on the red fluorescent coordinates (channel 1) using the green fluorescent coordinates (channel 2) as reference. The calculated registration map (named transform.npy) is saved in the /out_reg/ directory. The average distance between the beads’ centroid coordinates in each channel after correcting the CA is named the target registration error (TRE). Ideally, the calculated registration map should allow one to perfectly correct the CA between the two channels, thus retrieving a TRE ∼ 0 nm. However, this is often not the case (**5**). PyF2F calculates the TRE using the images of beads located in /test/ and saves the results (e.g. TRE.txt) in the /out_test/ directory.
23. Firstly, PyF2F subtracts the fluorescent signal corresponding to the extracellular background noise. Secondly, PyF2F calculates the background noise corresponding to the uneven cytoplasmic fluorescence signal at the edge of the cell using a median filter (a detailed description of why this step is important can be found in reference **48**). Finally, PyF2F subtracts the median filtered image from the image where the extracellular background had been subtracted. The final pre-processed images are saved at the /output/images/ directory under the names of “image_posXX.tif” for the extracellular background-subtracted image and “imageMD_posXX.tif” for the extracellular and intracellular background-subtracted images (“posXX” indicates the image number). We recommend using a rolling ball diameter slightly larger than the cell size, and a median diameter equal to twice the equivalent in pixels of the diffraction limit of the microscope. For our microscopy setup, we recommend using a rolling ball radius equal to 70 pixels and a median diameter of 11 pixels.
24. PyF2F uses trackpy (**49**) to localise independently the 2D-centroid coordinates of spots found in channel 1 and channel 2. Due to the experimental setup of PICT, the fluorescent signal of isolated and in-focus Sla2-RFP-FKBP platforms are recognized as diffraction-limited spots and are found at the cell contour (**7, 10, 48**). Channel 1 and channel 2 detected spots are saved at the /output/spots/ directory under the name of “detected_spot_posXX_W1*”* and “detected_spot_posXX_W2*”*, respectively. We recommend using a spot detection diameter of twice the diffraction limit of the microscope and a maximum separation distance between 2 to 3 pixels to link the detected spots in the two-channel into spot-pairs (“true” colocalising spots). Only linked spot-pairs are selected for posterior analysis. For our microscopy setup, we recommend using a spot detection diameter equal to 11 pixels.
25. The CA correction is applied to the channel 1 *xy* coordinates (files named “detected_spot_posXX_W1” at the /output/spots/ directory). The corrected coordinates are saved at the same location under the name of “detected_spot_posXX_W1_warped”.
26. The centroid coordinates are detected automatically and, because of the noisy nature of the imaging, potential false positives may contaminate the dataset. The filtering steps are meant to refine the dataset and reject as many spurious centroid localisations as possible. These steps worked well for our imaging setup and should be taken as an example that must be adapted to different acquisition setups. A stronger excitation, or a brighter fluorophore, would improve the localisation precision and make the dataset less noisy. Less refining would thus be needed. The filtering steps are the following:

● Close-to-cell contour spot-pairs: due to PICT experimental setup, detected spot-pairs must be located at the cell contour of the cell’s equatorial plane. PyF2F segments the cells by using a modified version of YeastSpotter (**46**). YeastSpotter uses a pre-trained Mask-CNN model to segment yeast cells imaged by fluorescence microscopy. We recommend using a distance-to-cell-contour cut-off equivalent to twice the diffraction limit in pixels, plus two pixels. For our microscopy setup, we recommend using a distance-to-cell-contour cut-off equal to 13 pixels. Moreover, PyF2F filters out overlapping spot-pairs based on a maximum closest-neighbour distance. We recommend using a maximum closest-neighbour distance equal to twice the diffraction limit minus one pixel. For our microscopy setup, we recommend using a maximum closest-neighbour distance equal to 10 pixels.
● High-quality spot-pairs: at this point, we assume that the majority of the selected spot-pairs are true positives (high-quality spot-pairs) and share similar properties in brightness and shape. Therefore, the majority of the spots will cluster in a two-dimensional space defined by the second momentum of brightness and the eccentricity for each RFP and mNG spot-pair. The clustering spots are identified with a 2D binned kernel density estimate (KDE). We recommend setting a threshold of 0.5 to select the spot-pairs that have a probability of 50% or higher to be found in the population (the 50% denser spot-pair population based on eccentricity and second momentum). The selected spot-pairs are those clustering in both channels.
● In-focus spot-pairs: the distribution of the pixel intensity in each spot is evaluated with a 2D Gaussian fitting. PyF2F evaluates the quality of the interpolation (*R^2^*) and selects spot-pairs above the threshold set by the user. We recommend using a threshold of *R^2^ >* 0.35 to select in-focus spot-pairs in both the channel 1 and channel 2.
27. The distribution of the distances between RFP and mNG spots is in fact skewed to the right (unless the two fluorophores are very far compared to the localisation precision, in which case the distribution approximates a normal distribution) (**3, 4**). The Maximum Likelihood Estimation (MLE) used to determine the true separation between the fluorophores is sensitive to outliers in a skewed distribution (**3, 4**). The rejection is thus based on the assumption that false negatives in the rejection of noisy data should retrieve a better outcome. PyF2F proceeds with a bootstrap method to iteratively reject outliers until one third of the dataset has been sampled for rejection (starting from the largest distances, which are those defining the tail of the distribution where outliers, if present, are more problematic). We recommend sampling 1⁄3 of the right-tailed area of the distance distribution. PyF2F selects the iteration that maximises the log-likelihood of the Rician distribution for the estimates of true distance (µ) and standard deviation (σ).

## Acknowledgement

We are grateful to Stephan Niekamp for technical support in the bead’s sample preparation and image registration steps of the protocol. We also thank Radovan Dojčilović, Marko Kaksonen and Baldo Oliva for helpful discussions. This work was supported by the Spanish funding agency (PID2021-127773NB-I00 funded by MICIU/AEI/10.13039/501100011033 / FEDER/UE, PRE 2019-088514 funded by MICIU/AEI /10.13039/501100011033 and by “FSE invierte en tu futuro”; the Unidad de Excelencia Maria de Maeztu (CEX2018-000792-M funded by MICIU/AEI/10.13039/501100011033); and the Human Frontiers Science Program (RGP0017/2020).

## Notes

https://github.com/GallegoLab/PyF2F

### Competing Interest Statement

The authors have declared no competing interest.

